# The enemy from within: a prophage of *Roseburia intestinalis* systematically turns lytic in the mouse gut, driving bacterial adaptation by CRISPR spacer acquisition

**DOI:** 10.1101/575076

**Authors:** Jeffrey K. Cornuault, Elisabeth Moncaut, Valentin Loux, Aurélie Mathieu, Harry Sokol, Marie-Agnès Petit, Marianne De Paepe

## Abstract

Despite an overall stability in time of the human gut microbiota at the phylum level, strong temporal variations in species abundance have been observed. We are far from a clear understanding of what promotes or disrupts the stability of microbiome communities. Environmental factors, like food or antibiotic use, modify the gut microbiota composition, but their overall impacts remain relatively low. Phages, the viruses that infect bacteria, might constitute important factors explaining temporal variations in species abundance. Gut bacteria harbour numerous prophages, or dormant viruses. A breakdown of prophage dormancy can evolve through the selection of ultravirulent phage mutants, potentially leading to important bacterial death. Whether such phenomenon occurs in the mammal’s microbiota has been largely unexplored.

Here we studied temperate phage-bacteria coevolution in gnotoxenic mice colonised with *Escherichia coli* and *Roseburia intestinalis*, a dominant symbiont of the human gut microbiota. We show that *R. intestinalis* harbors two active prophages, Jekyll and Shimadzu, and observed the systematic evolution of ultravirulent Shimadzu phage mutants, leading to a collapse of *R. intestinalis* population. In a second step, phage infection drove the fast evolution of host phage-resistance mainly through phage-derived spacer acquisition in a Clustered Regularly Interspaced Short Palindromic Repeats (CRISPR) array. Alternatively, phage resistance was conferred by a prophage originating from an ultravirulent phage with a restored ability to lysogenize.

Our results demonstrate that prophages are the potential source of ultravirulent phages that can successfully infect most of the susceptible bacteria. This suggests that prophages can play important roles in the short-term temporal variations observed in the composition of the gut microbiota.

## Introduction

The mammal’s gut microbiota is one of the densest microbial communities on earth. In Humans, its composition is globally conserved among healthy adults (Qin et al. 2010, Arumugam et al. 2011, Consortium 2013), and abnormal compositions, called dysbioses, are correlated with numerous human diseases (Backhed et al. 2004, Larsen et al. 2010, Marchesi et al. 2016). Yet the factors influencing the composition of the gut microbiota in health and disease are far from being entirely understood. For example, in a study on ∼1,000 healthy adults, the combined effect of 69 covariates (food, drug, anthropometric factors, etc.) could explain only 16.4% of the variation in microbiota composition (Falony et al. 2016). The impact of host genetics seems to be even lower (Rothschild et al. 2018). In addition, despite overall stability in term of presence or absence of main bacterial genera, strong variations in the relative abundance of species are observed on short time scale, even in healthy individuals submitted to the same diet (Koenig et al. 2011, Flores et al. 2014, Thaiss et al. 2014, Voigt et al. 2015, Sarker et al. 2017).

Bacteriophages, or phages for short, the viruses that infect bacteria, have been suspected to impact intestinal microbiota composition. In spite of clear evidence that phage-mediated selection plays an important role in bacterial strain diversification (reviewed in (Scanlan 2017)), there is as yet little proof that they are important determinants of the microbiota composition (reviewed in (Mills et al. 2013, De Paepe et al. 2014, Manrique et al. 2017)). Some metagenomics studies showed correlations between an increase in specific phages with reductions of particular bacterial taxa (Reyes et al. 2013, Waller et al. 2014). Yet, in most animal experiments using well-defined phage-bacteria pairs, phage-mediated bacterial mortality in the gut environment was limited to a fraction of susceptible bacteria, and was not sufficient to drive the selection of phage-resistant bacteria (Chibani-Chennoufi et al. 2004, Weiss et al. 2009, Maura et al. 2012), suggesting that intestinal physiology complicates phage multiplication (Brussow 2013). In line with this idea, the phage to bacteria ratio is much lower in the gastrointestinal tract than in other environments, suggesting less lytic replication (Silveira and Rohwer 2016). These different pieces of evidence might explain why the study of phages in the gut microbiota lagged behind the study of bacteria, and that phages are still often forgotten in microbiome studies.

Temperate phages, in particular, are generally considered to cause low bacterial mortality. They encode molecular mechanisms enabling them either to reproduce through lytic cycles upon infection, or to lysogenize bacteria, i.e. to establish in a stable dormant state in the infected cell. The dormant phage is called a prophage, the bacterial host a lysogen, and it is estimated that about 80% of bacteria are lysogens in the mammal’s gut (Touchon et al. 2016, Cornuault et al. 2018, Kim 2018). In line with this estimation, virions from temperate phages are dominant among intestinal viruses (Reyes et al. 2010, Minot et al. 2011, Knowles et al. 2016). Prophages are generally considered to be beneficial to their bacterial hosts, in particular in the gastrointestinal tract (Manrique et al. 2017, Mahony et al. 2018). The first advantage of lysogeny is superinfection inhibition, i.e. the protection from infection by other phages of the same “immunity” type. Inhibition of the superinfecting phage occurs by the same mechanism that establishes the dormancy of the resident prophage (Berngruber et al. 2010). Secondly, prophages can bring new functions to their host, favouring colonization, growth or stress resistance (reviewed in (Bondy-Denomy and Davidson 2014)). Finally, lysogeny can provide an advantage to the bacterial host by releasing infectious virions that are able to kill susceptible bacterial competitors (Bossi et al. 2003, Brown et al. 2006). Yet, in the gut, this advantage is limited by the frequent lysogenization of susceptible bacteria (Duerkop et al. 2012, De Paepe et al. 2016), as well as by the supposed rarity of susceptible competitors. Therefore, to date, the role of temperate phages in intestinal microbial communities is mostly studied from the perspective of horizontal gene transfer.

Yet some prophages can constitute a major threat for their hosts. In several microbial environments, important induction of prophages has been observed, resulting in the death of a significant part of the bacterial population (Selva et al. 2009, Brum et al. 2016, De Paepe et al. 2016). In addition, prophages of strains used in large-scale industrial fermentations are potential sources of virulent phage mutants infective for the host strain, the so-called ultravirulent mutants (Lucchini et al. 1999, Shimizu-Kadota et al. 2000, Durmaz et al. 2008). However, to the best of our knowledge, even though lysogeny is widespread among intestinal bacteria (Bobay et al. 2014, Touchon et al. 2016, Kim 2018), massive killing of bacteria by a temperate phage has never been documented in intestinal environments. This might result from a potential limited ability of phages to multiply in the gut environment as previously discussed, but also from a surprisingly paucity of studies of temperate phage-bacteria co-evolution in such an important microbial ecosystem (Scanlan, 2017).

The complexity of the gut microbiota prevents any exhaustive characterization of all the virus-bacteria interactions, even with high throughput metagenomic studies. A particular roadblock is the difficulty to predict the bacterial host of the phages detected, in particular for phages with no relatives among reference sequenced viruses (reviewed in (Edwards et al. 2016)). In particular, no phage infecting *Lachnospiracea*, the most abundant bacterial family of the human gut microbiota (Falony et al. 2016), has been deposited in public databases. Here we followed the co-evolution of a *Lachnospiracea* strain, *Roseburia intestinalis* L1-82, and its prophages in gnotoxenic mice. The *Roseburia* genus is of particular interest not only because of its abundance in the gut microbiota, but also because of its important production of short chain fatty acids in the human gut, including butyrate, which have a critical role in the regulation of host immune responses (Rios-Covian et al. 2016). In addition, this genus is depleted in several human diseases, suggesting it might be important for human health (Tamanai-Shacoori et al. 2017). In our study, mice were also colonized with *Escherichia coli*, a sub-dominant member of the human gut microbiota whose abundance increases in several human diseases (Willing et al. 2010). We show that ultravirulent mutants of *R. intestinalis* phage Shimadzu systematically invade this simplified gut ecosystem, and lyse the large majority of susceptible bacteria. Phage mediated bacterial killing drives the fast evolution of host phage-resistance through phage-derived spacer acquisition in a Clustered Regularly Interspaced Short Palindromic Repeats (CRISPR) array. Our results illustrate that prophages are not always dormant, and can play important roles in the short-term temporal variations observed in the composition of the microbiota.

## Results

### *Roseburia intestinalis* L1-82 harbors two active prophages

The sequencing and complete assembly of *R. intestinalis* L1-82 genome revealed the presence of two complete prophages, that we named Shimadzu and Jekyll (Fig. 1A). The production of virions by these prophages was determined by sequencing encapsidated DNA, i.e. DNA protected from DNAse in a culture supernatant. Resulting reads predominantly aligned on the two identified prophages (60% and 19.4% of total reads respectively), indicating that both prophages produce virions. Of note, read coverage was highly uneven on the Jekyll prophage, suggesting that besides the full size phage genome, a shorter version is also encapsidated (Fig. S1A). Phage genomes were annotated using RAST (Aziz et al. 2008), HHPRED (Zimmermann et al. 2018) and Phagonaute (Delattre et al. 2016) as previously described (Cornuault et al. 2018). Interestingly, the Shimadzu integration site is located within its integrase gene *int*, encoding a large serine recombinase. Consequently, two different integrases are produced during the lytic cycle and during lysogeny (Fig. S1B). The two forms differ by their last 22 amino acids, which might modify the stability of the protein, as described in mycophages (Broussard et al. 2013).

**Figure 1:**
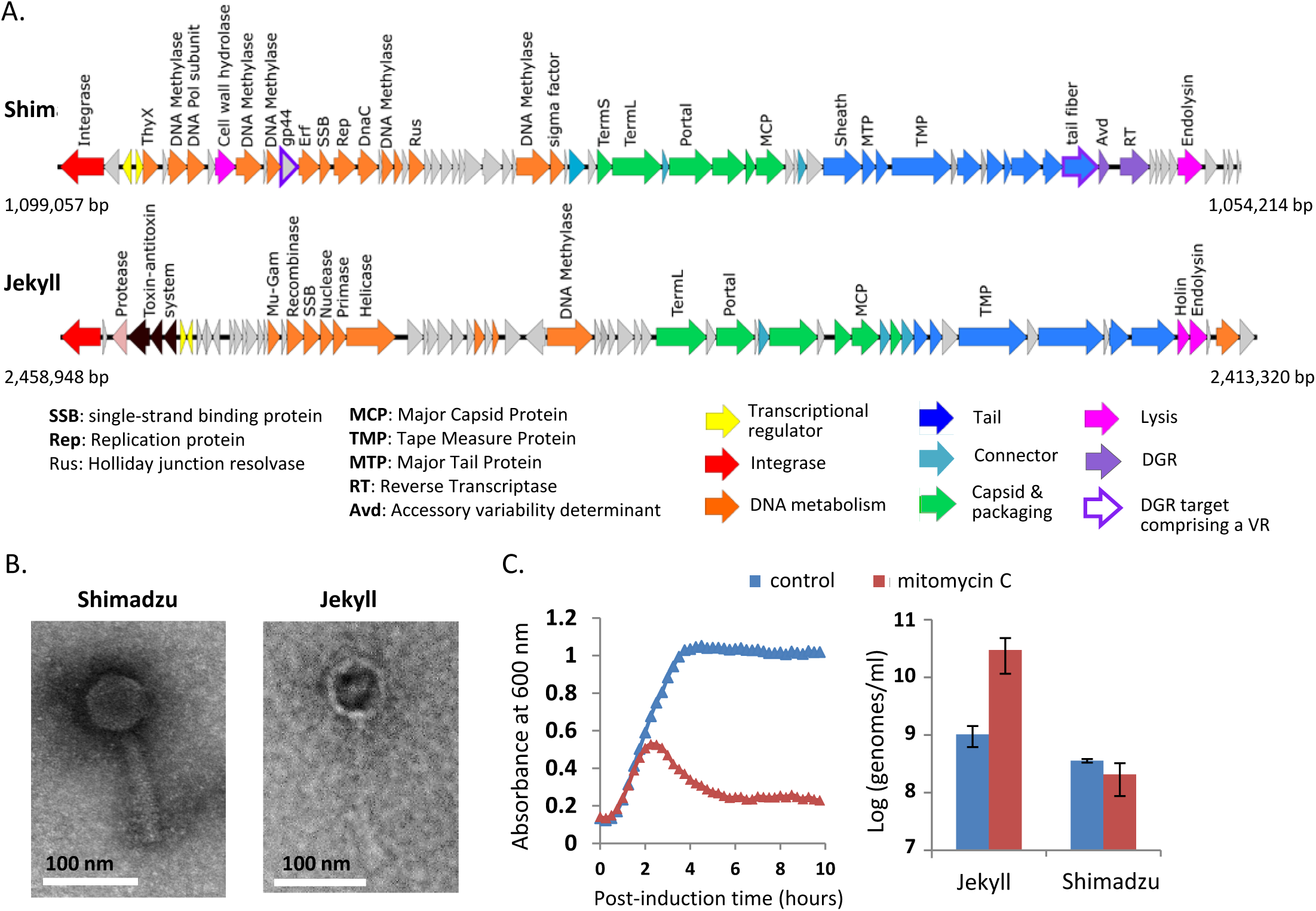
*Roseburia intestinalis* L1-82 prophages. **A)** Genetic maps of Shimadzu and Jekyll prophages. Positions in *R. intestinalis* L1-82 genome are indicated. **B)** Transmission electron microscopy images of Shimadzu and Jekyll virions. **C)** Phage genome concentrations in bacterial cultures with or without mitomycin C (0.5 µg/ml). Mean ± standard deviation of three measures.

The Jekyll genome had no significant nucleotide similarity with any viral genome of the NCBI RefSeq genome database, indicating that it corresponds to a new viral clade. The Virfam classification tool, that uses virion protein remote homology and gene synteny (Lopes et al. 2014), clustered Jekyll with phages infecting *Bacilli* species, such as lactobacillus phage LLH. The Shimadzu genome presented homology with the *Faecalibacterium prausnitzii* phage Lagaffe (Fig. S1C), and grouped within a badly-resolved Virfam cluster comprising *E. coli* lambda phage. Electronic microscopy showed that Jekyll is a *Siphoviridae*, and Shimadzu a *Myoviridae* (Fig. 1B), in line with genomic predictions. Quantitative PCR on encapsidated viral DNA indicated that the virion concentration in saturated culture supernatants was 3.10^8^ and 8.10^8^ virions/mL for Shimadzu and Jekyll respectively. Interestingly, Jekyll, but not Shimadzu, was induced by mytomicin C, a genotoxic agent causing DNA damage, a general signal for prophage derepression (Fig. 1C).

Similarly to *F. prausnitzii* phage Lagaffe, Shimadzu encodes a Diversity Generating Retroelement (DGR) (Fig. S2). DGRs generate variability in target genes through a reverse transcriptase (RT)-mediated mechanism that introduces nucleotide substitutions at defined locations, called variable repeats (VR) (reviewed in (Guo et al. 2014)). VR are homologous to a template repeat (TR), located directly upstream the reverse transcriptase, and used as a template to generate variation. As sometimes observed (Wu et al. 2018), two VR could be detected in Shimadzu: the first one, VR_1_, is located in a predicted tail gene located just before the reverse transcriptase, a commonly observed position, and the second, VR_2_, is located in *orf43*, of unknown function (Fig. 1A and S2B). Analysis of sequence reads obtained from virions collected from *in vitro* culture supernatants indicated that both VR are successfully modified by the DGR: 2 % of reads aligning to the VRs had 10 to 15 mismatches typical of DGR mediated mutagenesis, whereas only 0.1% of reads outside this region exhibited 1 to 2 mismatches.

### *R. intestinalis* and Shimadzu populations exhibit important temporal variations in mice

We then examined free phage and bacterial populations in germ-free mice colonized with *R. intestinalis* L1-82 and either *E. coli* LF82 (experiments 1 and 2) or *E. coli* MG1655 (experiment 3). Each experiment comprised 5 to 8 mice housed in two different cages placed in the same isolator. Phage and bacterial populations were monitored by quantitative PCR and/or microscopy in faeces for 33 days (Fig. S3). *R. intestinalis* became rapidly dominant over *E. coli*, reaching 96 ± 2.5% of the population after 5 days of colonization (Fig. 2A). However, in each cage, a significant collapse of *R. intestinalis* was observed in the majority of mice after 16 to 25 days of colonization (Fig. 2A). Since mice were kept in a constant and closed environment, we hypothesized that one of the two *R. intestinalis* L1-82 prophages could be responsible for this change in bacterial composition. During the first 10 days of colonization, the concentration of Jekyll and Shimadzu virions in mouse faeces was comparable to that observed *in vitro*, around 5 x 10^8^ virions/g of faeces (Fig. 2B). Afterward, Jekyll free phage concentration remained constant but Shimadzu concentration suddenly increased by more than a thousand fold, reaching up to 10^12^ virions/g of faeces after 13 to 25 days of colonization, before decreasing and stabilizing around 4 x 10^10^ virions/g (Fig. 2B). Similar results were observed in mice colonized with the two different *E. coli* strains, ruling out potential effects of *E. coli* LF82 virulence genes on phage production.

**Figure 2:**
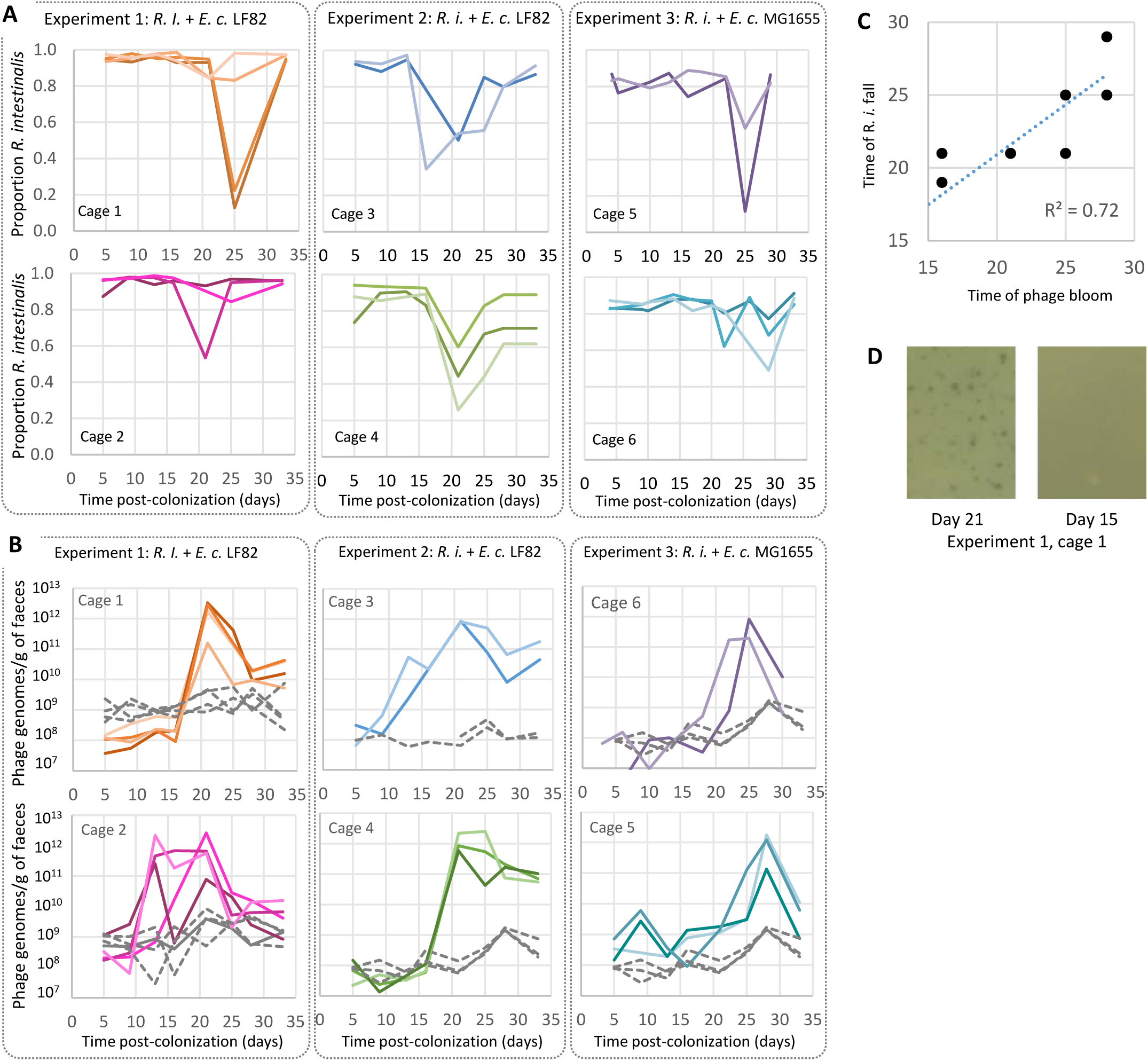
Evolution of *R. intestinalis* and phage populations with time in mouse faeces. Each panel corresponds to a different cage, hosting two to four mice. In each experiment, two cages (1 & 2, 3 & 4, or 5 &6) were placed in the same isolator. Mice from cages 1 to 4 were colonized with the LF82 strain of *E. coli*, whereas mice in cages 5 and 6 were colonized with the *E. coli* MG1655 strain. **A)** Evolution of the proportion of *R. intestinalis* over total bacteria. These proportions were estimated by quantitative PCR in experiments 1 and 2, and by microscopy in experiment 3. **B)** Evolution of the concentration of Shimadzu (solid line) and Jekyll (dashed lines) free-phage genomes. **C)** Time of *R. intestinalis* population collapse as a function of time of Shimadzu bloom. **D)** After 21 days of colonization, lysis plaques could be obtained from mouse faeces (left panel), but not from faeces collected at earlier time points (right panel).

### Shimadzu multiplication is responsible for *R. intestinalis* mortality

Shimadzu free phage concentration was maximal shortly before the fall of concentration of *R. intestinalis* (Fig. 2C, R_2_ = 0.72), strongly suggesting phage-related bacterial mortality. Yet, as discussed earlier, prophages provide resistance to superinfection, i.e. lysogenic bacteria are immune to infection by a phage that is present as a prophage in its genome. The sudden multiplication of Shimadzu could thus result either from the invasion of ultravirulent mutants able to infect lysogenic cells, or from a sudden environmental signal triggering a massive prophage induction. In favour of the first hypothesis, Shimadzu lysis plaques on *R. intestinalis* L1-82 lawn were obtained from mouse faeces collected after 21 days of colonization (Fig. 2D). These lysis plaques, by definition, were formed by ultravirulent mutants. Of note, the number of plaque forming units (PFU) obtained corresponded to 10^2^ to 10^6^ PFU per gram of faeces, far from the 10^12^ virions estimated by quantitative PCR. Quantification of three phage stocks revealed a 100-fold difference between genome copy numbers determined by qPCR and PFU, suggesting that most virions are uninfectious in these conditions, but not entirely explaining the difference observed in faeces between PFU and virions. This difference could result either from damage of virions during extraction or from genetic heterogeneity in the population of Shimadzu in mouse faeces, with only few phage genotypes being able to infect *R. intestinalis* L1-82 *in vitro*. Nevertheless, the isolation of phages on L1-82 indicated that Shimadzu ultravirulent mutants were present in mice, most probably explaining bacterial mortality.

### The bloom of Shimadzu free phage is associated with few dominant mutations

To identify the mutations responsible of the ultravirulent phenotype, we sequenced five phages, isolated either from experiment 1 (Shivir1) or from experiment 2 (Shivir2, 3 and 4) after 21 days of colonisation, and Shivir5 from extra cage 7 of experiment 2, not fully presented in this report, and isolated after 33 days of colonisation. Each phage genome contained between 34 and 45 mutations, but only four loci were mutated in the five isolates: the two VRs targeted by the DGR (∼10-15 mutations per VR), a region comprising two inversely oriented transcriptional regulators, named immunity region, and finally *orf26*, of unknown function (Fig. 3A). To know if these mutation hotspots represented a general trend rather than biases due to phage isolation, we sequenced total phage DNA (virome) isolated from mouse faeces harvested after 21 days of colonisation. Virome 1 corresponds to a pool of faeces from the 8 mice of experiment 1, whereas virome 2 corresponds to a single mouse of experiment 2. In both viromes, mutations in the immunity region, in VR_1_ and in VR_2_ were respectively present in ∼70%, ∼65% and 47% of aligning reads. No other locus was mutated in more than 20% of covering reads (Fig. 3A&B). For example, the mutation in *orf26* was present in only 5 % of aligning reads in virome 1, and even less in virome 2, indicating that this mutation is not essential for Shimadzu multiplication in mice. Non-synonymous mutations in all five phage isolates (Fig. 3C) suggests mutations in *orf26* are nevertheless necessary to form lysis plaques *in vitro*, and might explain the discrepancy between PFU and phage copy numbers in faeces discussed above.

**Figure 3:**
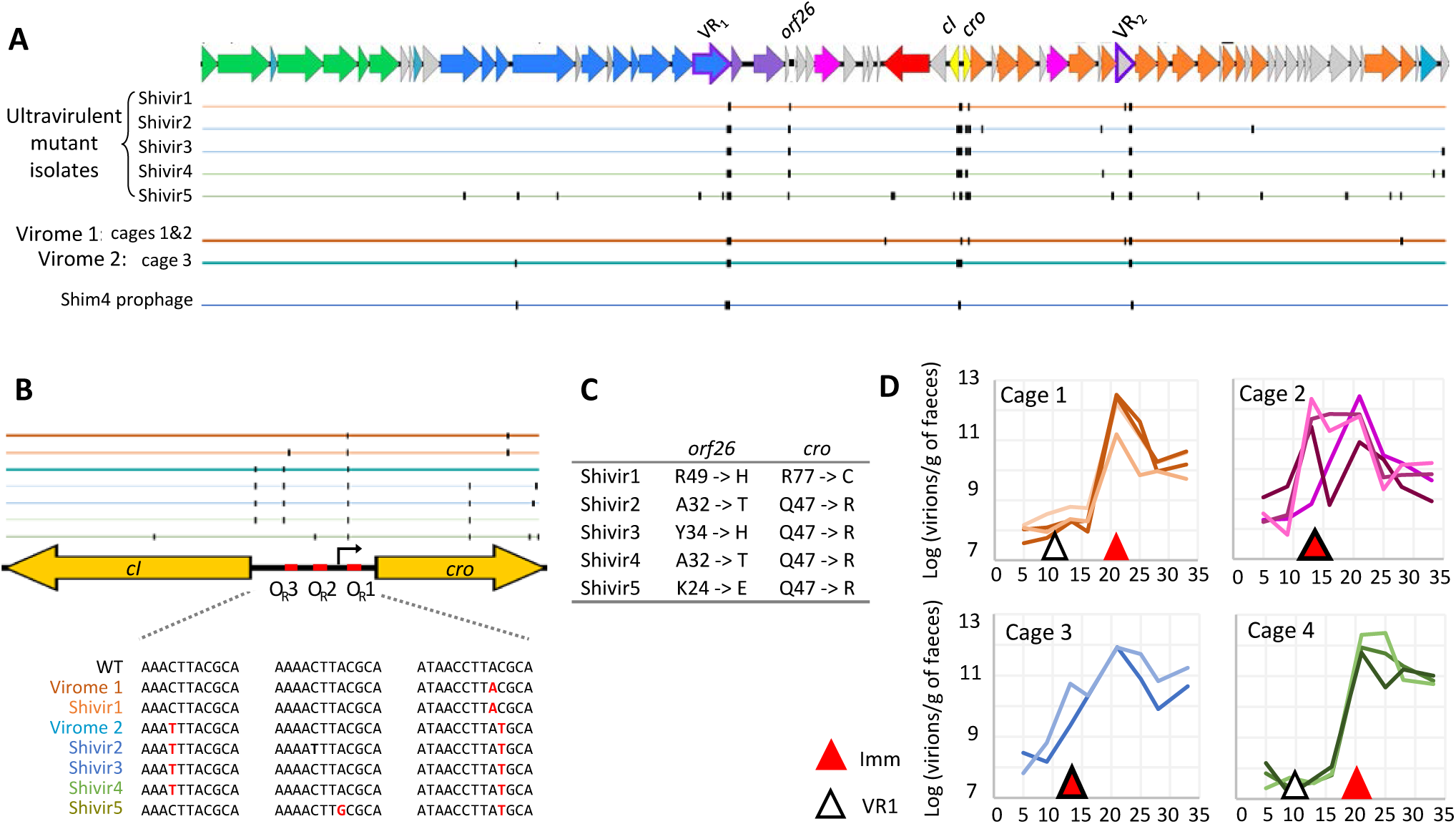
Mutations of ultravirulent Shimadzu phages. **A)** Representation of mutation positions in virome 1 (all mice from experiment 1) and 2 (one mice from cage 3 in experiment 2), and in ultravirulent phage isolates. Color of the line indicates the cage of isolation. **B)** Upper panel: Mutation positions in the immunity region of Shimadzu, comprising the repressor genes *cI* and *cro* and the intergenic regions with three operators. Lower panel: mutations in the operators. **C)** Amino acid mutated in *orf26* and *cro* in ultravirulent isolates. **D)** Time of first detection of mutations in the operators and in the VR1 in total viral DNA from mouse faeces.

In the two viromes, VR_1_ and VR_2_ variants comprised one to three dominant alleles, with the exception of the VR_2_ in virome 1, in which the most abundant allele represented only 3% of reads (Table S1). All ultravirulent isolates from experiment 2 had the same VR_1_ allele, which corresponds to the second most dominant allele in the virome 2 (Table S1). By contrast, all isolates carried different VR_2_ alleles, and differed from the most abundant ones in the corresponding virome (Table S1). This reflects stronger selection of VR_1_ alleles as compared to VR_2_, probably resulting from a higher importance of the VR_1_ containing tail protein for infection.

The mutated region comprising two inversely oriented transcriptional regulators is most probably the immunity region, that controls prophage maintenance and resistance to superinfection. Indeed, similarly to the immunity region of the model phage lambda, the intergenic region comprises three almost identical repeats, known as operators (O_R_) in lambda phage, and a putative promoter identified with the dedicated tool DBTBS (Sierro et al. 2008) (Fig 3B). In phage λ, the two transcriptional regulators *cI* and *cro* are under mutually exclusive expression: during lysogeny, O_Rs_ are fixed by CI repressor, preventing the expression of *cro* and the downstream lytic cycle genes, and vice versa during the lytic cycle. Upon superinfection, CI repressors will also bind the O_R_ of the incoming phage, preventing the expression of its lysis genes, and conferring immunity. All five ultravirulent phages have a mutation in *cro* and in O_R_1 (Fig. 3B&C). These mutations might diminish or even abolish the binding of CI repressor produced by the resident prophage, enabling the incoming phage to multiply in lysogens. Mutations in O_R_2 or O_R_3 were also observed in some isolates, probably accentuating this phenotype.

### Dynamics of mutation selection

To gain insights into the respective role of the mutations described above in Shimadzu multiplication and phage bloom, we then examined their timing of selection (Fig 3C). Since dominant VR_2_ alleles in virome 1 represented only 3% of sequences and that each phage isolate had a different allele, we concluded that the VR_2_ containing gene might be less essential for Shimadzu multiplication, and we focused on the VR_1_ and the immunity region. Important VR_1_ diversification was detected before the onset of the phage bloom in two cages, indicating the selection of mutant alleles prior to phage bloom. This selection suggests that even prior to the bloom, the production of free phage not only results from spontaneous prophage induction but also from an infection process. This infection probably concerns only a small subpopulation of bacteria since neither an increase in free phage titers, nor a bacterial decrease, could be detected at that time. Such VR_1_ allele selection could occur if the ancestral tail fiber gene is not permitting efficient infection. We hypothesized that the infected bacterial population could consist of bacteria having lost spontaneously the Shimadzu prophage. Screening of clones isolated either from an *in vitro* culture or from mice after 13 days of colonisation revealed in the two cases a proportion close to 5.10^-3^ cured bacteria, a proportion confirmed by qPCR. We can speculate that these cured bacteria are the susceptible hosts enabling VR_1_ allele selection. O_Rs_ mutations became dominant concomitantly with the phage bloom in all mice, definitely associating the major phage multiplication peak with the evolution of ultravirulence.

### Ultravirulent phages drive the fast selection of resistant bacteria

Despite the systematic and very important multiplication of Shimadzu in all mice, *R. intestinalis* concentration almost recovered its initial level after 30 days of colonization, and in some mice the bacterial population was barely affected by the ultravirulent phages (Fig. 2). These observations hint at the rapid emergence of resistant bacteria. To investigate this phenomenon, a total of 300 *R. intestinalis* clones were isolated after 16, 19, 25, 28 or 33 days of colonization in experiment 2. Ninety-six of them were grown in the presence or absence of ultravirulent phage Shivir5, which showed the best lysis ability of bacterial liquid cultures. For two additional Shivir mutants, a smaller scale analysis (on 10 bacterial clones) by plaque assay revealed an identical susceptibility pattern. The proportion of resistant clones dramatically increased after the phage bloom (Fig. 4A), explaining the recovery of *R. intestinalis* population in mice.

**Figure 4:**
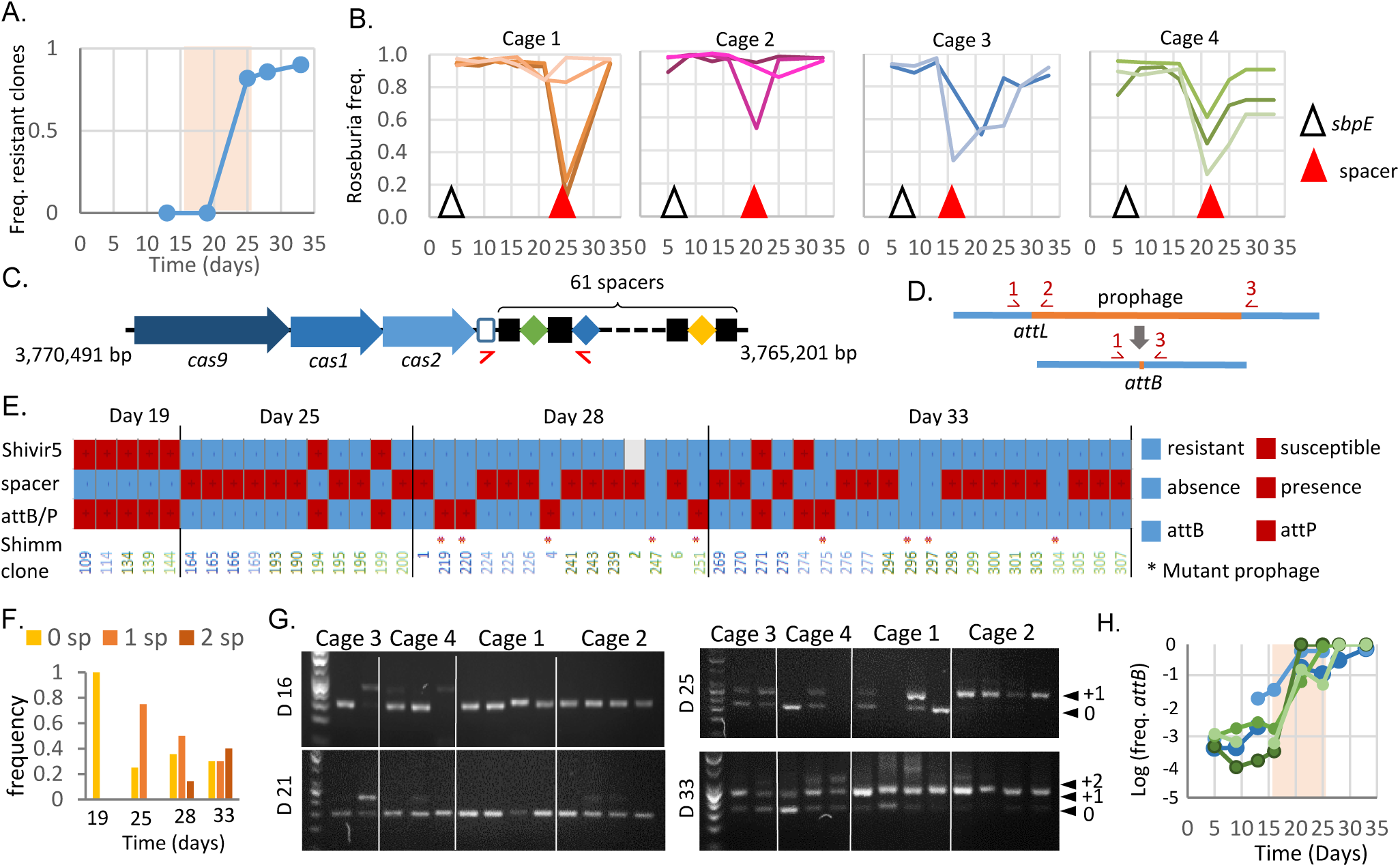
Phage resistance is associated to spacer acquisition or lysogenization by a prophage with a new immunity type. **A)** Proportion of bacterial clones resistant to Shimadzu with respect to isolation time from cages 3 & 4. The pale pink region indicates the Shimadzu bloom. **B)** Time of first detection of mutations in *sbpE* and spacer acquisition in total bacterial DNA from mouse faeces. **C)** Type II-C CRISPR-cas system of *R. intestinalis* L1-82 and position of primers used for detection of spacer acquisition. Positions in the bacterial genome are indicated. **D)** Schematic representation of Shimadzu *attL* and *attB* sites, and their detection by PCR. **E)** For 52 isolated clones, are indicated: Shivir5 resistance (upper line), spacer acquisition (middle line), presence of *attB* or *attL* sites (lower line), and presence of a Shim prophage (*). The color of clone number corresponds to the mouse from which it has been isolated, as in Fig. 2. Sequenced clones correspond to numbers 1, 2, 4 and 6. Clones isolated at earlier time points were all identical to those of day 19 and therefore not included in the figure. **F)** Frequency of clones with 0, 1 or 2 new spacers as a function of isolation time. **G)** Spacer acquisition in total *R. intestinalis* bacteria from faeces at four different time points (day 16, day 21, day 25 and day 33). DNA ladder is a 50 pb scale. Smaller PCR fragments correspond to the wild-type size (indicated by “0”), the acquisition of 1 spacer to “+1” and 2 spacers to “+2”. **H)** Population level of Shimadzu cured *R. intestinalis*, in mice from different cage. Cage is color coded as in B panel.

### Resistance is mostly related to the acquisition of phage-directed CRISPR-Cas spacer

Bacteria have developed an incredible number of mechanisms to counter phage infections (reviewed in (van Houte et al. 2016)). To get insight into the mechanisms involved, 4 resistant clones of *R. intestinalis* isolated after 28 days of colonization were entirely sequenced. Each of them contained between 38 and 54 mutations, but only gene *sbpE* (RiL182_04007), presenting homology to a *Bacillus subtilis* extracellular solute-binding protein gene, was mutated in all clones. Yet this mutation was dominant in the mouse gut early on, after only 5 to 7 days of colonization (Fig. 4B), and long before the Shimadzu bloom, suggesting that *sbpE* mutations are associated to adaptation to the mouse gastro-intestinal tract rather than to phage resistance.

Sequencing also revealed in three of the four isolates the insertion of a 30 base pair sequence homologous to the Shimadzu genome (Table S2), in a Clustered Regularly Interspaced Short Palindromic Repeats (CRISPR) array, of type II-C (Fig. 4C). CRISPR and their associated genes, CRISPR-Cas immune systems, provide phage resistance by integrating short phage-derived sequences (spacers), generally at the leader end of the CRISPR array, similarly to what we observed. Type II-C CRISPR-Cas systems function by cleaving invading DNA homologous to spacer sequences. Spacer acquisition was then investigated in the clones tested for Shivir5 susceptibility: more than 80% of the resistant clones had acquired a new spacer, but none of the susceptible clones (Fig. 4C&E), and the proportion of clones with two additional spacers increased with time (Fig. 4F). Sequencing of newly acquired spacers in 10 other clones showed that they all possess one or two Shimadzu-derived spacers, but two clones also possess a spacer that targets the *R. intestinalis* genome (Table S2). The already described cost associated with auto-immunity of CRISPR-cas type II systems was apparent because these two clones displayed altered growth, especially on plates. Finally, spacer sequencing revealed the presence in the same mouse of clones with different spacers, revealing several spacer acquisitions in the same mouse.

To confirm that bacterial resistance through CRISPR-Cas activity is general and does not result from biases in the isolated clones, we monitored spacer acquisition on total bacterial DNA from mouse faeces (Fig. 4G). In each cage, important spacer acquisition was first evidenced at the onset of bacterial recovery (Fig. 4B). In addition, the number of acquired spacer increased with time, exactly as in isolated clones (Fig. 4F). Altogether, these results indicate that CRISPR-Cas activity is the main *R. intestinalis* resistance mechanism against ultravirulent Shimadzu in the mouse gut.

### Spacer acquisition is associated with the loss of Shimadzu prophage

Full genome sequencing of the three *R. intestinalis* clones with an additional spacer also revealed their loss of the Shimadzu prophage. Amplification by PCR of either *attB*, the site formed by prophage excision, or *attL*, a site formed by prophage presence (Fig. 4D), revealed the absence of Shimadzu in all 32 clones with a new spacer (Fig. 4E). To evaluate the representativeness of the isolated clones, we determined by quantitative PCR the evolution with time of the global proportion of faecal bacteria with intact *attB* site. This proportion remained close to that estimated by screening of individual clone, 5.10^-3^, for the first two weeks of colonization (Fig. 4H). Yet, after phage bloom, the proportion of bacteria with an intact *attB* site dramatically increased in all tested mice: Shimadzu-derived spacer acquisition most probably selected cured bacteria because of the important cost associated with autoimmunity.

### Resistance is also mediated by a prophage with a modified immunity region

By contrast, the fourth sequenced clone (clone 4 in Fig. 4E), had no additional spacer but a second Shimadzu prophage, inserted in a tRNA at position 3,542,165 of the genome. Sequence analysis revealed that some prophage positions were mutated in 50% of aligning reads, suggesting that the second prophage, named Shim4 (for Shimadzu prophage with modified Immunity region located in clone 4), was mutated at these positions (Fig. 3A). In particular, two different alleles of both VRs could be identified, and the VR_1_ mutant allele was identical to that of the ultravirulent Shivir phages isolated from the same experiment (Table S1). No mutation in *orf26* was identified, reinforcing the idea that such mutations are not essential for infection *in vivo*. In the immunity region, the same three point mutations observed in corresponding virome 2 were detected, but the *cro* gene and its promoter were inverted (Fig. 5A). This inversion might lead to an increased expression of CI repressor, enabling to restore the resistance to superinfection of the lysogen. To investigate the generality of this phenomenon, we looked for the presence of such Shim prophage with an inversion of *cro*, in the eight other clones resistant to Shivir5 infection but that had not acquired an additional spacer. We could amplify by PCR the Shimadzu immunity region in all isolates, even in clones with an intact *attB* sites, indicating the presence of a prophage at another genomic position than in the ancestral L1-82. An inversion of *cro* gene was evidenced in all eight isolates (Fig. 4E). These bacterial isolates still produced virions, albeit at a level much lower than the ancestral L1-82 strain or other lysogens isolated from mice, comforting the hypothesis that they produce more CI repressor in the prophage state than the L1-82 (Fig. 5B and S4).

**Figure 5.**
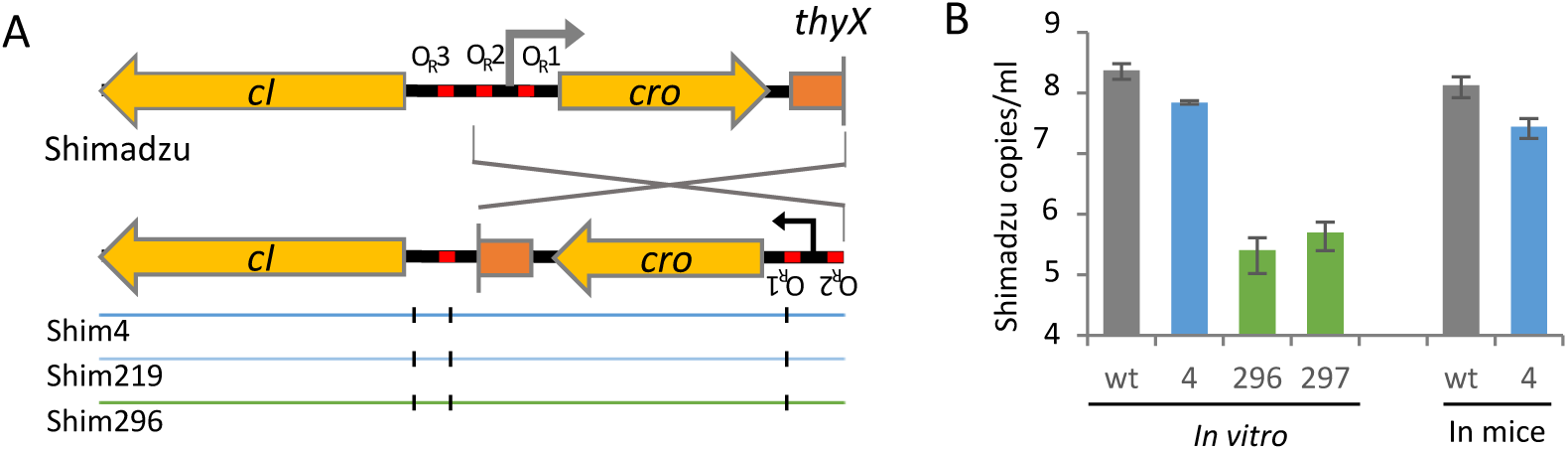
Prophage Shim has an inverted *cro* region. **A)** Zoom on the immunity region, showing the *cro* inversion in Shim prophage and the 3 point mutations. **B)** Spontaneous Shimadzu virion production in clones 4, 296, 297 and in L1-82 ancestral clone, *in vitro* of after 3 days of mice colonization. Clone 4 is a dilysogen harbouring Shimadzu and Shim, the other clones only contain a Shim copy. Mean ± s.d of three measures.

Altogether, ultravirulent phage resistance of the 40 clones tested can be explained either by spacer acquisition or by lysogenization by a phage with a new immunity type. For clarity, a summary sketch of the results is proposed in Fig. 6.

**Figure 6.**
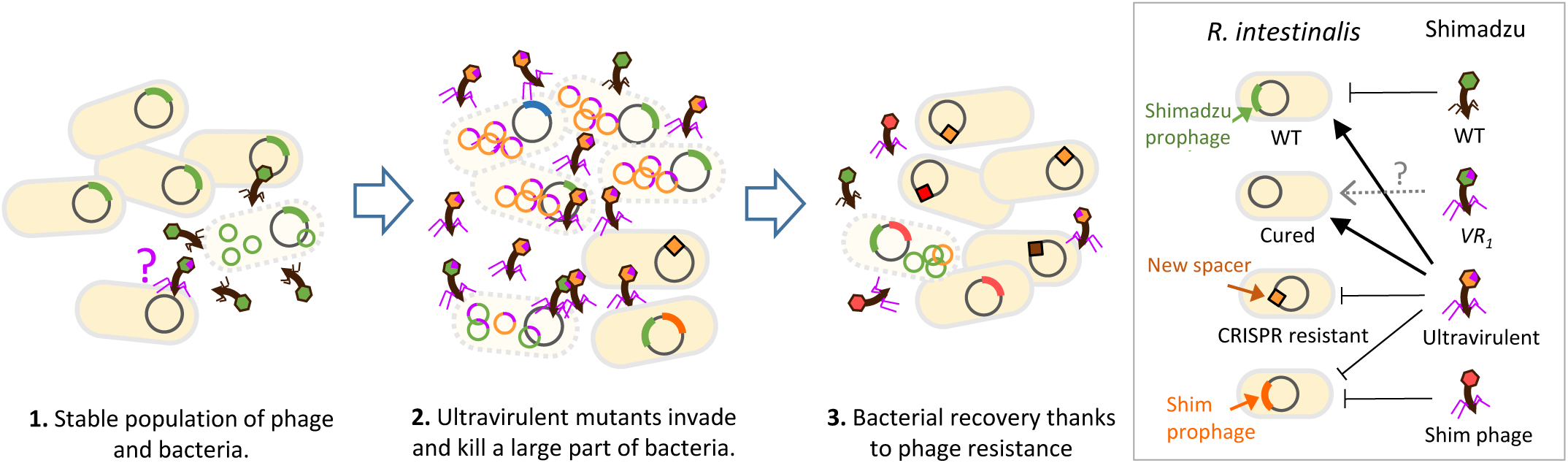
Summary sketch of the results. In a first step, stable populations of free phage and bacteria are observed. Yet, complex dynamics might already occur, as 0.1% of bacteria are cured of the Shimadzu prophage, and phages with mutant VR1 alleles are selected. The second step corresponds to the free phage bloom: ultravirulent Shivir mutants invade and multiply on bacteria. Phages with a restored lysogenisation ability (Shim phages) are generated. In the third step, resistant bacteria invade, thanks to CRISPR spacer acquisition and to lysogenization by a new immunity type prophage.

## Discussion

The prophage state is generally considered a dormant state that gives little opportunity for viral evolution, since viruses mutate much faster during lytic replication than during lysogenic cycles (Sanjuan et al. 2010). This might explain why the coevolution between temperate phages and bacteria have been much less studied than between virulent phages and bacteria (Koskella and Brockhurst 2014, Scanlan 2017). Yet, theory predicts that ultravirulent mutants with mutations in the operator that diminishes the binding of the repressor should invade the population when transmission is high, i.e. when the benefit of superinfection outweighs the cost of virulence (Berngruber et al. 2010, Berngruber et al. 2013). Here, we followed the coevolution between *R. intestinalis* L1-82 and its prophages for 33 days in the mouse gut, and demonstrate that Shimadzu ultravirulent mutants systematically invade, leading to major phage-mediated mortality of *R. intestinalis*. In addition, in agreement with theory (Berngruber et al. 2010), we observed in some cases later invasion of compensatory mutations that restore lysogenization ability and provide resistance to the ultravirulent mutants. Since the large majority of bacteria of the human microbiota harbours prophages (Touchon et al. 2016, Kim 2018), this kind of dynamics might regularly occur, leading to massive phage-induced bacterial mortality, and participating in the variations in microbiota composition.

Even though predicted by theory, evolution of ultravirulence was considered to only rarely arise in nature, and, to the best of our knowledge, had only been observed in industrial setups, where large bacterial population size favours the appearance of mutants (Shimizu-Kadota et al. 2000, Capra et al. 2011, Mercanti et al. 2011). Based on results with lambda phage, the best-known example of lysogeny regulation, this rarity was proposed to stem from the fact that a double mutant would need to be generated, and the probability of acquiring two simultaneous mutations is very small during lysogeny (Lenski and May 1994, Koch 2007). Because of this presumed rarity, the evolution of ultravirulence is rarely, if ever, considered when referring to lysogeny in host-associated microbiota. Similarly to what was shown with lambda, all ultravirulent Shimadzu mutants had at least two mutations in the immunity region, in operators O_R_1 and O_R_3. Yet, when a single strain dominates at high concentration, as in gnotoxenic mice, our results show that ultravirulent mutants with several simultaneously acquired mutations can be generated. Single-strain stability, i.e. the presence of a single dominant strain, was observed for numerous species of the gut microbiota (Truong et al. 2017), and in particular for *Roseburia* (Moss et al. 2017), suggesting that the environmental conditions necessary for ultravirulence evolution should be present in natural gut ecosystems. Most probably, some specific phage genetic features are also required. In particular, whether the phage DGR system and/or the presence of spontaneously cured bacteria facilitated ultravirulence evolution is a very interesting open question that deserve further investigation in a future work.

Here phage infection resulted in the fast selection of resistant bacteria in the mouse gut, contrary to what is generally observed *in vivo* (Chibani-Chennoufi et al. 2004, Maura et al. 2012, Reyes et al. 2013, Nale et al. 2016). This absence of genetic resistance selection was interpreted as a result of physiological bacterial resistance to phage infection in the mouse intestine, resulting from its complex structure and slow bacterial growth (Miedzybrodzki et al. 2012, Brussow 2013, Berngruber et al. 2015, Maslov and Sneppen 2015). In our experiments, almost all bacterial clones isolated after the emergence of ultravirulent phages were resistant, indicating very efficient phage-mediated selection and thereby particularly efficient multiplication of Shimadzu in the mouse gastrointestinal tract. To the best of our knowledge, our work constitutes the first examination of phage-bacteria interactions with a dominant member of the gut microbiota, and such outcome might be not so rare with this kind of bacteria. Ultravirulent prophage mutants might be responsible from some of the unexplained variations in bacterial populations in animal experiments.

Despite this very efficient phage infection, bacterial population did not drop so much, and this is due to rapidly acquired resistance by CRISPR-Cas immunity. CRISPR-Cas systems are widely distributed in prokaryotes, but only a small proportion of these systems have been identified to be active in bacteria. Here we demonstrate very good efficiency of the minimal type II-C CRISPR-Cas system of *R. intestinalis*. Considering that among strains acquiring spacers only those cured of Shimadzu were conserved, and that cured bacteria were about 10^7^ per mice, we can estimate a rate of spacer acquisition above 10^-7^, similar to that of the best active CRISPR-Cas systems (reviewed in (McGinn and Marraffini 2019)). This high efficiency might be driven by the presence of defective Shimadzu particles, suggested by the discrepancy between Shimadzu PFU and the number of free-phage determined by qPCR. Indeed, there is evidence that spacer acquisition occurs most efficiently during the infection by defective viral particles (Hynes et al. 2014). Yet, interestingly, phage-derived spacer acquisition did not lead to phage extinction. This could result from a high phage mutation rate, enabling to generate escape mutants, but also from the presence of lysogens with an additional Shim-like prophage, that continuously produce free phage.

Interestingly, because the bacteria harbour a prophage closely related to the invading phage, spacer acquisition is highly detrimental, and leads to a kind of suicide. In the case of ultravirulent phage mutants, CRISPR–Cas does not act as a true immunity system, but rather functions as an abortive infection system leading to infected cell death. Luckily for *R. intestinalis*, a subpopulation of spontaneously cured bacteria, around 5.10^-3^, is sufficient for phage-derived spacer acquisition and recovery of the bacterial population. However, contrary to Shimadzu, most prophages are almost impossible to cure (our own experience). In some cases, this might be related to the presence of toxin-antitoxin genes, as in the Jekyll prophage of this study, that stabilize the presence of prophages through post-segregational killing (Lehnherr et al. 1993, Romero et al. 2009, Goeders and Van Melderen 2014, Niu et al. 2015). In consequence, such subpopulation of cured bacteria does not always exist, probably precluding type II CRISPR-Cas efficiency against superinfection by ultravirulent mutants.

In the present study, constitutive resistance to Shivir conferred by phage receptor mutations was not observed. This is surprising in the light of studies showing that phage receptor mutations tend to dominate, especially when phage populations are high (Westra et al., 2015). The fitness cost of phage receptor mutations, in the case of Shimadzu, may be even higher than the one associated to CRISPR-Cas activity. But an alternative explanation could be that such receptor mutants may also be systematically countered by Shimadzu escape mutants, thanks to the DGR-mediated high variability of its tail gene which possibly encodes the receptor binding protein.

Finally, our study illustrates how a prophage can be an important source of mortality in bacterial populations. Therefore, prophages could be compared to “genetic time bombs”, meaning that lysogenic bacteria live under the constant threat of superinfection by an ultravirulent mutant. Since phages are known to alter competition among bacterial strains or species (Bohannan and Lenski 2000, Joo et al. 2006, Koskella et al. 2012, Li et al. 2017), such evolution could have major effects on the composition of an ecosystem. In particular, *Roseburia* species are very important in the human gut microbiota, notably due to its important butyrate production, a metabolite with critical role in the regulation of host immune responses (Rios-Covian et al. 2016). Phage-mediated killing of *Roseburia* bacteria might therefore participate to the unexplained variations in microbiota composition observed even in defined communities (Koenig et al. 2011, Flores et al. 2014, Thaiss et al. 2014, Voigt et al. 2015, Sarker et al. 2017), and, in certain conditions, lead to important changes and a rupture of intestinal homeostasis.

## Material and methods

### Bacterial cultures

*R. intestinalis* L1-82 (DSM 14610), was grown at 37°C in LYBHI medium (BHI medium (Merck) supplemented with 0.5% yeast extract (Difco), cellobiose at 0.5 mg/ml (Sigma), maltose at 0.5 mg/ml (Sigma) and cysteine at 0.5 mg/ml (Sigma) in an anaerobic chamber filled with 90% N2, 5% CO2 and 5% H2. *Escherichia coli* LF82 was grown in LB medium (Difco) at 37°C. *E. coli* was grown on LB agar plates (1.5% agar). *R. intestinalis* was plated on LYBHI agar plates (LYBHI medium with 1.5% agar and hemin at 0.5 %), supplemented with 2µg/mL of ciprofloxacine to suppress *E. coli* growth when necessary.

### Sequencing of encapsidated DNA in L1-82 culture supernatant

500 ml of a L1-82 culture grown overnight were centrifuged at 5,200 g for 15 min at 4 °C. Supernatant was recovered and centrifuged at 5,200g for 30 min at 4 °C. This step was repeated with 1 h of centrifugation and the samples were concentrated with PEG and NaCl as described below. Phage pellet was resuspended in 2 ml of SM buffer (100 mM NaCl, 8 mM MgSO4, 50 mM Tris pH 7.5 (Sigma) and treated with 0.25 μg of RNAse A (Sigma) and Dnase I (Sigma) at 37 °C for 1 h. Phage DNA was then extracted using Promega kit Wizard® DNA Clean-up system. Sequencing was performed with the Ion proton sequencing technology at the INRA Metagenopolis platform. Alignment of reads was performed using bowtie2 (-N 1 –L 32) and then visualized with Tablet using default parameters (Milne, 2013).

### Mitomycin C induction

Overnight cultures were diluted 100-fold and grown at 37°C until absorbance at 600nm was between 0.2 and 0.4. 135 µL of culture was distributed in a 96-well plate (Cellstar, 96 Well Cell Culture Plate, U-bottom) and 15µL of H_2_O or Mitomycin C (Sigma)) was added at the appropriate dilution. Plate were sealed (AMPLIseal^TM^ Greiner Bio-one) in anaerobic condition and incubated at 37°C in a TECAN SUNRISE. Absorbance at 600nm was measured every 15 min for 10 hours. Phage genome counts in supernatant were determined 6 hours after induction by quantitative PCR as described below.

### Animals and experimental design

Germfree 5 to 8 week-old C3H/HeN mice (female) from the germfree rodent breeding facilities of Anaxem-Micalis (INRA, Jouy-en-Josas, France) were kept in flexible-film isolators (Getinge-La Calhène, Vendôme, France) in standard cages (2 to 4 mice/cage) with sterile wood shavings as animal bedding. Mice were given free access to autoclaved tap water and to a standard diet, R03-40 (Scientific Animal food and Engineering, Augy, France), sterilized by gamma irradiation. Isolators were maintained under controlled conditions of light (12h), temperature (20 to 22°C) and humidity (45 to 55%). To obtain gnotobiotic *E. coli*/*R. intestinalis*-diassociated mice, mice were orally inoculated with 100µL of a saturated *E. coli* LF82 culture (2.5×10^9^ CFU). 48 hours later, 10^10^ CFU of *R. intestinalis* in 200µL were administered by intragastric gavage of mice pretreated with sodium bicarbonate (0.2 M, 0.1mL by intragastric gavage, 10 min of wait before inoculation of bacteria). All procedures were carried out according to European Community Rules of Animal Care and with authorization 1234-2015101315238694 from French Veterinary Services.

### Phage extraction and isolation from faeces

Fresh or frozen faeces thawed on ice for 10 min were diluted 40-fold in cold PBS. Resuspended faeces were kept on ice for 5 min with regular agitation prior to centrifugation for 10 min at 5,251 g at 4°C. Supernatants were recovered and filtered through a 0.22 μm filter (PALL Corporation Acrodisc PF syringe filter). For subsequent quantification by PCR, virions were precipitated with PEG 8000 (Sigma) at a final concentration of 10% and NaCl (Sigma) at a final concentration of 1M. After overnight incubation at 4°C, phage particles were harvested by centrifugation (5,251 g for 1 hour at 4°C with a swinging rotor) and resuspended in 100 μl of SM buffer (100 mM NaCl, 8 mM MgSO4, 50 mM Tris pH 7.5 (Sigma)). Samples were treated with 10 U of Turbo DNase (Ambion) for 1 hour at 37 °C in Turbo DNase buffer and then incubated at 95°C for 30 min in 0.2 ml PCR tubes for enzyme inactivation and capsid disruption. For phage isolation, the fecal filtrates were mixed with 150 µL of an exponentially growing *R. intestinalis* culture (Optical density at 600nm comprised between 0.2 and 0.5) in an anaerobic chamber. 1 ml of melted Top BHI agarose (LYBHI supplemented with 0.25% agarose and 0.1% cysteine (Sigma)) at 37°C was added and the mix was immediately poured on 6 cm Petri Dishes containing LYBHI agar. Plates were then incubated overnight at 37°C in the anaerobic chamber. A single lysis plaque was then mixed with an exponentially growing bacterial culture and top agar containing and poured on an LYBHI Agar plate as described above. After 16 hours of incubation at 37°C, lysis was confluent, and 2 ml of SM buffer were poured on the top of the plates and incubated at 4°C for two hours. The phage supernatant was then recovered and filtered at 0.2 µm.

### Bacterial DNA extraction from faeces

Bacterial pellets obtained during free phage extraction from faeces were resuspended in 500 µL of LB (Difco). The suspension was centrifuged 1 min at 500 g to remove debris. Supernatant was recovered and mixed with 250 µL of lysis buffer (200mM NaCl/20mM EDTA, 5% SDS), 250 µL of phenol:choloform:isoamyl alcohol (25:24:1; [pH 8.0], Sigma aldrich) and half of a tube of silica beads (100µM MP Bio, Lysing matrix B). Bacteria were lysed using Fast-prep MPBio (5.5, 3×30s, 5 min between each cycle). Samples were then centrifuged (13,000 g, 3min, 20°c) and the aqueous phase recovered. After addition of 400 µL of chloroform:isoamyl alcohol (24:1), the mix was vortexed vigorously (10s) and centrifuged (13,000 g, 3min, 20°C). DNA was precipitated with 2 volumes of ethanol and 0.3M of sodium acetate and resuspended in 100 µL of water.

### Phage and bacteria quantification by qPCR

qPCR on 100-fold diluted samples was performed using the Takyon ROX SYBR Mastermix blue dTTP kit (Eurogentec) and the StepOnePlus real time PCR system (Applied Biosystem). Phage and bacteria amounts were determined using specific primer pairs (Table S3). Numbers of copies in standards were calculated using DNA quantification (Qbit) and the strain genome size. The reaction mix was the following: 7.5 μl Takyon mix, 0.9 μl H_2_O, 0.3 μl of each primer (200 nM final concentration), 6 μl of DNA diluted in H_2_O. The PCR conditions were: (95°C 15s, 58°C 45s, 72°C 30s) 45 cycles, 72°C 5 min, followed by melting curves. Results were analysed using the StepOne Software 2.3.

### Isolation of *R. intestinalis* clones from mouse faeces

One or two faeces were recovered from mice, sealed immediately in screw cap micro tube, transferred to an anaerobic chamber (see bacterial culture), diluted in 0.5 mL of PBS 1X. 20µL of this bacterial suspension was then streaked on GHYBHI plates (brain-heart infusion medium supplemented with 0.5% yeast extract, 1.5% agar, cellobiose (1 mg/ml [Sigma]), maltose (1 mg/ml [Sigma]), cysteine (0.5 mg/ml [Sigma]), Hemin 0,5% and ciprofloxacin at 2 µg/mL). After 24 to 48h of growth at 37°C, isolated clones were grown 24h at 37°C in liquid LY-BHI medium, frozen with 20% of glycerol and kept at −80°C until analysis.

### Quantification of Shimadzu cured bacteria

In an anaerobic chamber, bacteria either from a growing culture (OD600 = 0.3) or from a frozen faeces were rapidly diluted in cold PBS to prevent reinfection and plated on LYBHI agar plates. Individual clones were used to inoculate two 96 well microplates per condition. After a 16 hours incubation at 37°C, PCR with either attB or attL primers (table S3) was realized on 1 µl of bacterial cultures, and the proportion of attB positive clones determined. For quantification by PCR, quantitative PCR was performed on bacterial DNA as described above with attB and sigA primers, and the proportion of cured bacteria determined as the attB over sigA copy numbers.

### Shimadzu susceptibility assay in 96 wells microplates

A 96 well microplate (Cellstar, 96 Well Cell Culture Plate, U-bottom) was inoculated with 96 different clones and duplicated for conservation at −80°C and subsequent tests. For each test, a microplate was used to inoculate a microplate filled with 200 µl of LYBHI. Overnight cultures were diluted 50-fold and grown at 37°C for two hours in 50 µl. 5 µl of a phage lysate at a concentration of 10^9^ pfu/ml was added and the microplate incubated for 30 mn at 37°C. Cultures were then diluted 4-fold in warm LYBHI and incubated at 37°C for 16 hours. Absorbance at 600nm was then measured in a TECAN SUNRISE, and compared to the absorbance of cultures grown in similar conditions but with no phage. Clones were considered susceptible to the phage if the absorbance at 600nm of the culture with phage was more than 2-fold lower than that of the culture without phage. The experiment was repeated two times for all clones and a third time for clones that gave incongruent results.

### Shimadzu susceptibility assay on agar plates

A 5 µl drop of phage suspension containing ∼10^5^ PFU was spotted on a lawn of *R. intestinalis* in top agarose as described above, and incubated at 37°C for 16 hours. Confluent lysis at the position of the drop indicated bacterial susceptibility, whereas 10 or less PFU indicated bacterial resistance.

### Bacterial numeration by microscopy

10-fold dilutions of freshly passed faeces in PBS were spread on an agarose pad on a microscope slide and imaged at 100x magnification by a microscope (Zeiss Axioskop2Plus; camera Zeiss axiocam ICc1), in phase contrast. *E. coli* and *R. intestinalis* cells were manually counted (Fig. S3). We analysed more than 50 cells on 3 different images for each condition. In parallel, *E. coli* CFU were numerated on LB agar plates. The concentration of *R. intestinalis* cell was obtained by multiplying the concentration of *E. coli* deduced from plating by the ratio of *R. intestinalis* on *E. coli* cells obtained by microscopy.

### Transmission electronic microscopy (TEM)

For Jekyll virion observation, 100 mL of *R. intestinalis* culture supplemented with 1µg/mL of mitomycin C were centrifuged and the supernatant filtered through a 0.2 μm filter, concentrated with PEG and 0.5M NaCl as described above and then purified by ultracentrifugation (20,000g, 2 h, 4 °C) on a 3 layers iodoxanol gradient (concentration of 0, 20% and 45% of iodoxanol). Virions were recovered at the interface between the 20% and the 45% layers. Shimadzu virions were extracted from mice faeces resuspended in PBS. After centrifugation for 2 min at 10,000g, the supernatant was filtered and then concentrated using an ultrafiltration device Amicon Ultra 2 mL with a cut-off of 100 000 Da (Millipore). For both phages, droplets of phage preparations were directly placed on Formvar carbon-coated grids for 5 min. The grids were stained with 1% uranyl acetate and then viewed for TEM using a HITACHI HT 7700 (Elexience, France) at 80 kV. Microphotographies were acquired with a charge-coupled device camera AMT.

### *R. intestinalis* L1-82 sequencing, assembly and annotation

Genomic DNA was extracted from a bacterial culture with a standard phenol:chloroform protocol preceded by a lysis step with lysozyme and SDS, as previously described (Fouet et al., 1990). Sequencing was realised by the Genotoul GIS, using the PacBio technology. Long reads were quality controlled whith NanoPlot v1.8.1 (De Coster et al. 2018). Sequencing produced 202,572 reads of N50 12,694 nucleotides for a total run of 1,214,051,359 bases. This corresponds to a sequencing depth of 270x. Assembly was performed using Canu 1.6 (Koren et al. 2017) with default parameters except genomeSize=4.5m and pacbio=raw. Two contigs were produced by the assemblers, one corresponding to the chromosome, the other of 21,201 nucleotides with a GC% of 0.01%. The second was considered spurious and removed from the analysis. A run of Illumina paired-end sequencing (5,827,730 paired-reads of 151 bp) was conducted to correct the remaining error. Reads were first mapped on the PacBio assembly using BWA 0.7.12-r1039 (Li and Durbin 2009), then the assembly was corrected using Pilon 1.22 (Walker et al. 2014). 290 corrections of the initial assembly were performed, mostly indels correction. A final single circular contig of 4 493 348 nucleotides was obtained, and its first nt was chosen at the shift of the genome GC skew, near the *dnaA* gene. The genome was then annotated with Prokka (Seemann 2014), and prophage genes and boundaries were manually edited. The new version of the genome has been deposited on the EBI database under accession number ERZ773328. On Shimadzu phage, promoters of *cI* and *cro* genes were searched with DBTBS (http://dbtbs.hgc.jp/).

### Detection of mutations in the sequenced bacterial isolates

Research of SNPs in bacterial genomes was made using the reference L1-82 genome, using reads generated by Illumina sequencing by GATC-Eurofin and analysed using the PATRIC web interface (https://patricbrc.org) (Wattam et al. 2017). Mapping was made using BWA-MEM and SNP were searched with freebayes with default parameters. Prophage insertion in clone 4 was first evidenced by the twice higher read coverage on the region corresponding to the Shimadzu prophage, following reads alignment using bowtie2 (-N 1 –L 32) and visualization with Tablet using default parameters (Milne 2013). The position of the second prophage was determined by recovering, at each prophage border, the sequence of 10 clipped reads. These “hybrid” reads were then aligned with BLASTn to the L1-82 genome, revealing the insertion site. Spacer insertion was detected as in other clones, as described below.

### Shivirs and virome sequencing

5 ml of ultravirulent phage stocks (concentration above 10^9^ PFU/ml) were concentrated with 10% PEG8000 and 0.5M NaCl. Alternatively, for virome sequencing, virions were extracted from two to three mouse faeces as described above for qPCR. In both cases, viral DNA was then extracted with a standard phenol:chloroform protocol. DNA was sequenced either by GATC-Eurofins with the Illumina technology (2×125 nt), or, in the case of virome 1, with IonTorrent (300 nt reads) in the INRA metagenopolis unit.

### Shivir mutation analysis

Raw reads were quality filtered using Trimmomatic (Bolger et al., 2014) in order to remove paired reads with bad qualities (with length < 100 bp, with remnant of Illumina adapters and with average quality values below Q20 on a sliding window of 4 nt). The filtering process yielded an average of 1 to 3 million paired-end reads (2 × 150 bp) per sample. Point mutations and small indel were detected by alignment of reads using bowtie2 (-N 1 –L 32) and visualized with Tablet using default parameters (Milne 2013). Since Shivir were multiplied on L1-82 strain that carry the ancestral Shimadzu prophage, ∼20% of reads corresponded to the wild-type Shimadzu sequence. Putative larger deletions or insertions were searched by assembling the filtered reads using SPAdes (Bankevich et al., 2012) with the meta option and increasing kmer values (-k 21,33,55,77,99,127). The resulting contig was then compared by BLASTn to the ancestral Shimadzu.

### Analysis of viromes

A first search of SNP was performed similarly to the Shivir analysis. The more thorough analysis of the VR regions was performed as follow. First, using Tablet, all reads covering the VR region of interest and 20 extra nt on each side (for 300 nt-long reads, virome 1), or at least one side (for 125 nt-long reads, virome 2) were downloaded. We reasoned that the various VR alleles in a given virome should group together in a phylogenetic tree. These reads were therefore used for a multi-alignment with MUSCLE (Edgar 2004), which was then trimmed with Gblocks, and given to PhyML on the LIRRM interface (Dereeper et al. 2008) to build a tree (default parameters, when more than 500 reads were present, two subpopulations were made and treated in parallel). Several clear flat branches per tree were obtained, corresponding to the main alleles of a given VR. The number of leafs of each such cluster was counted, and by substraction, the remaining reads were considered as singleton alleles. A read of each main cluster was then used to determine the amino-acid sequence of each VR allele. To determine allele proportions, a mapping of virome reads on each VR allele was performed with bowtie2, allowing 0 mismatch (-N 0 –L 20).

### Detection of spacer acquisition

Spacer acquisition was determined by a PCR test (primers in table S3). PCR products were observed by electrophoresis on a 2% agarose gel.

### Detection of mutations by Sanger sequencing

Sanger sequencing of the *R. intestinalis sbpE* and CRISPR loci was performed by GATC-Eurofin on DNA substrates generated by PCR amplification of bacterial DNA purified from mouse faeces with primers in table S3, using standard procedures. Sequences of the Shimadzu VR_1_ and immunity region were similarly obtained from PCR amplification products of viral faecal DNA. Manual inspection of sequencing traces allowed to detect peak mixtures, indicative of the presence of mutants in the population.

## Acknowledgements

We are very grateful to Anne Foussier, Fatima Joly and Aurélie Balvay from Anaxem facility for their help with animal handlings, to Christine Longin from the MIMA2 facilities (UMR 1313 GABI, INRA) for the TEM observations, and to the Migale platform (INRA) for the bio-informatics environment. JKC was funded by the Fondation pour la Recherche Médicale (FDT20170437017), and mouse experiments were funded by Association François Aupetit.

## Author contributions

JKC, EM and MDP performed the experiments. VL assembled and annotated the *R. I.* L1-82 genome. JKC, MAP and MDP performed the mutation analyses. AM, HS and EM developed protocols. JKC and MDP analyzed the results and wrote the manuscript, and all authors revised and approved the manuscript.

## Supplementary table S1

**Table S1.**
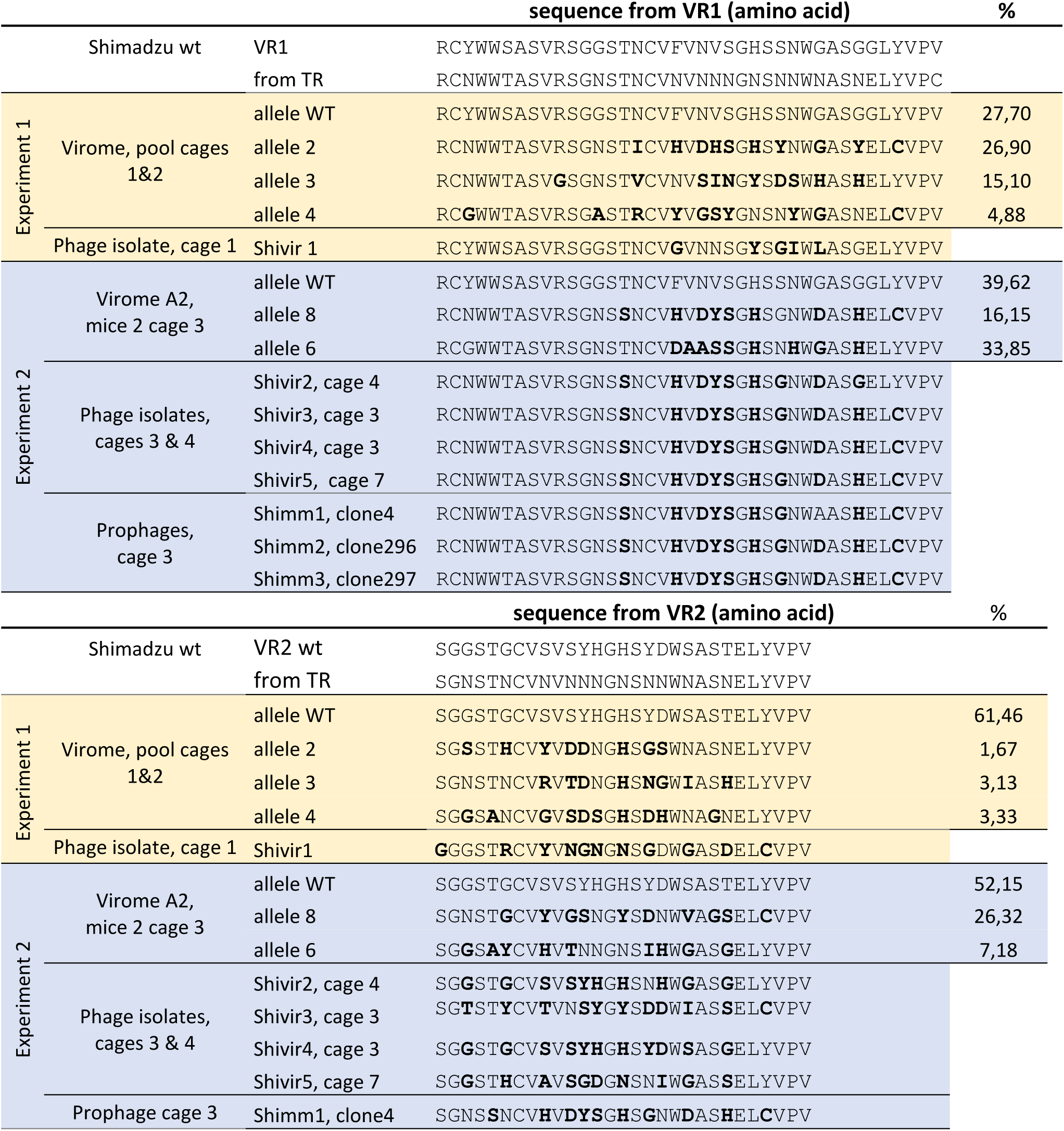
Amino acid sequences of the protein segment containing the DGR VRs, in viromes and in Shimadzu isolates. For the Shimadzu WT, two sequences are indicated: the first (VR_1_) corresponds to the actual protein, while the second correspond to the translation of the template repeat (TR). Color indicates amino acids modified relative to this TR sequence. For the viromes, the percentage (%) of each allele is indicated.

**Table S2.**
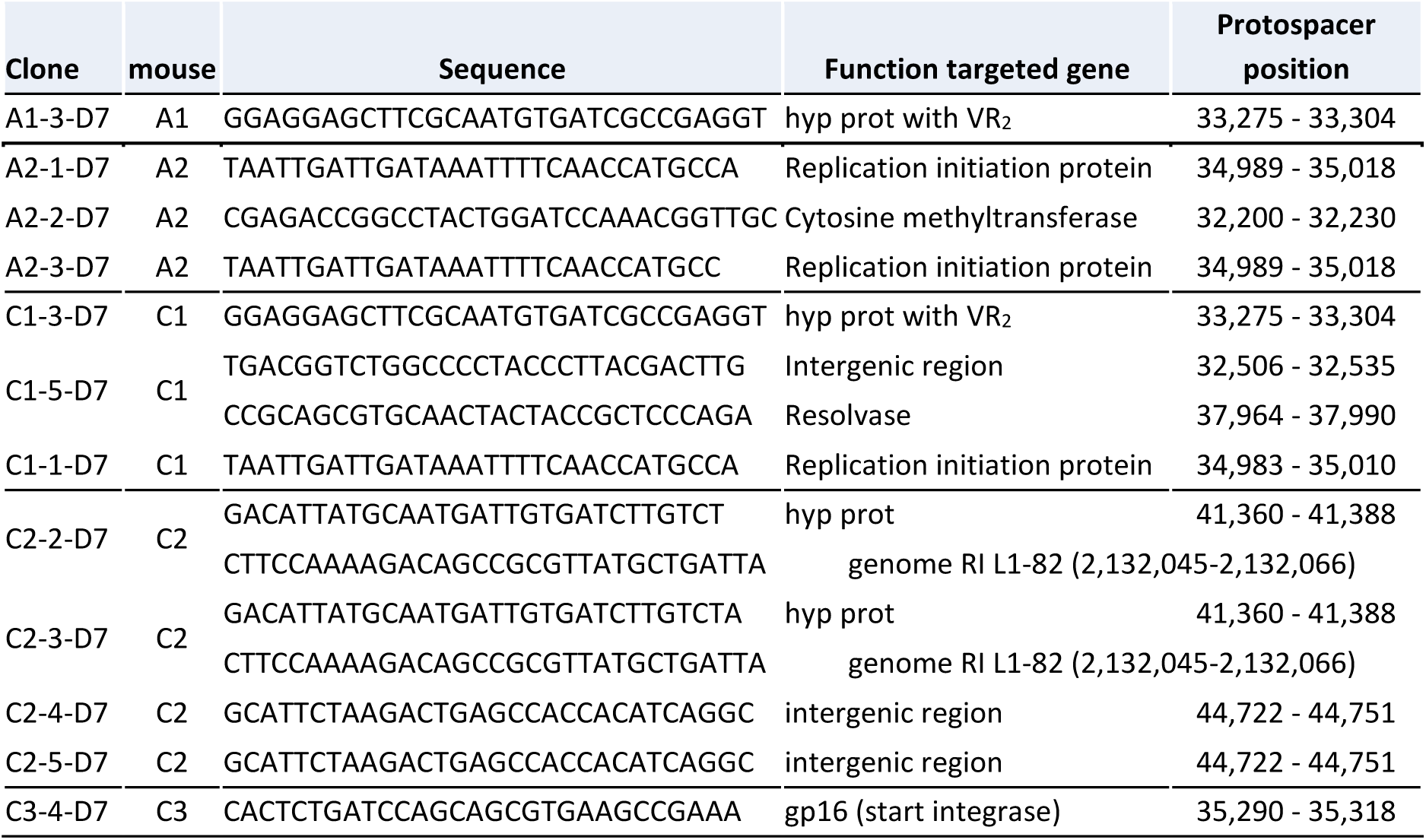
Sequences and corresponding protospacer positions of acquired spacers.

**Table S3:**
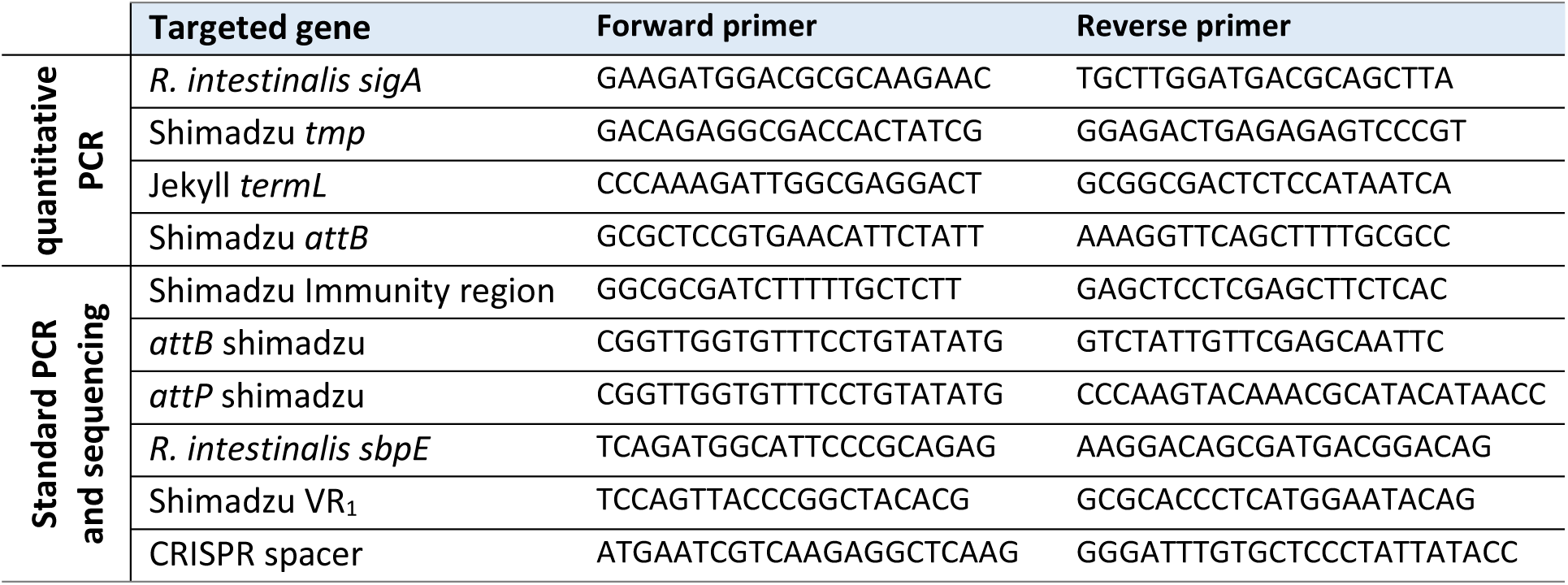
Oligonucleotides used in this study.

**Figure S1.**
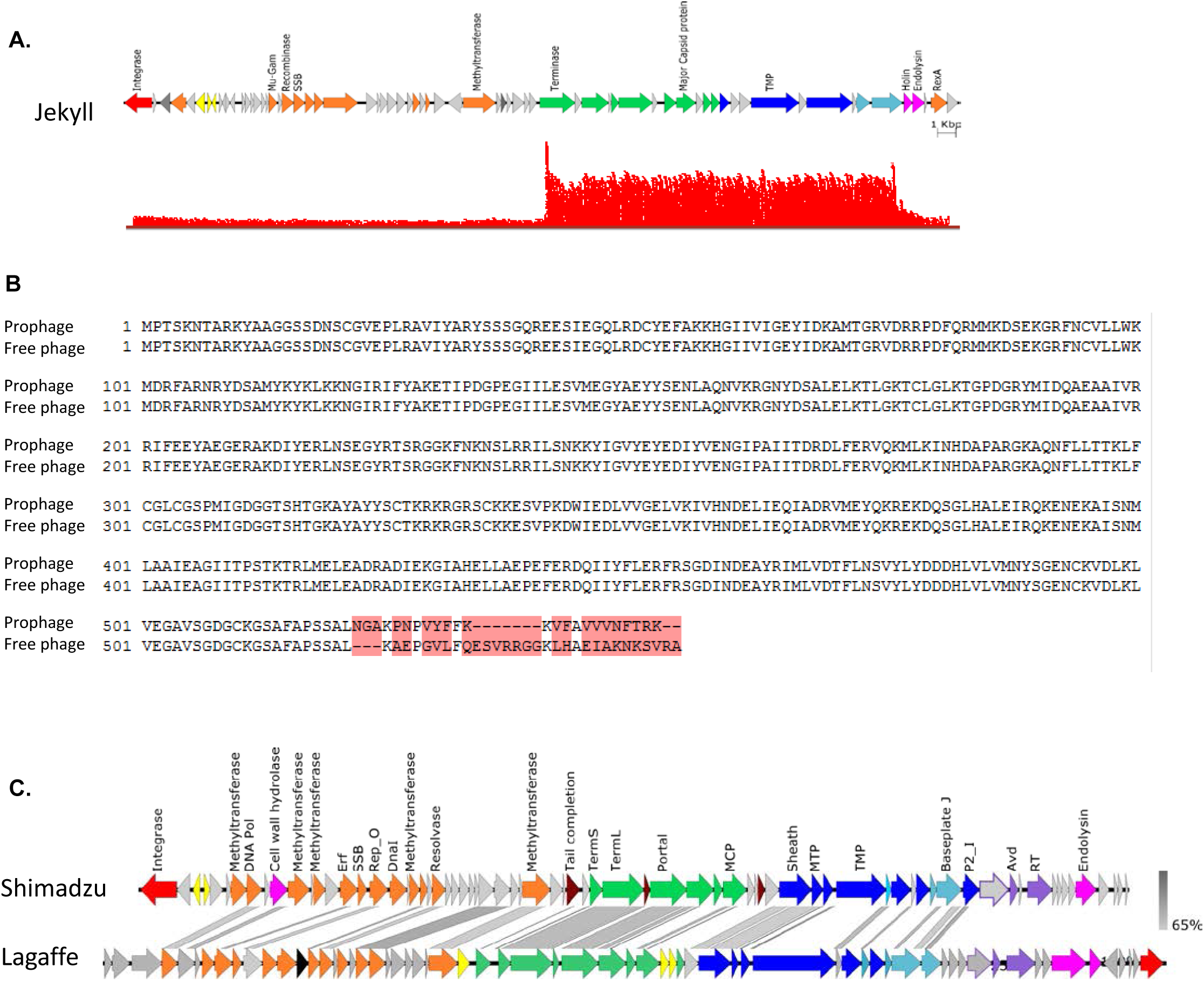
Peculiar genomic features of the two prophages. A) Read coverage of Jekyll prophage. The region comprising capsid genes present an 8-fold higher coverage than the rest of the genome. B) Alignment of the Shimadzu integrase sequence in the prophage (upper line) and free phage (lower line) forms. The last 20 amino acids are different. C) Shimadzu and Lagaffe (GenBank MG711461) genomic comparison. Blast was made using BlastN. The two phages share 60% of their proteins and overall good synteny.

**Figure S2.**
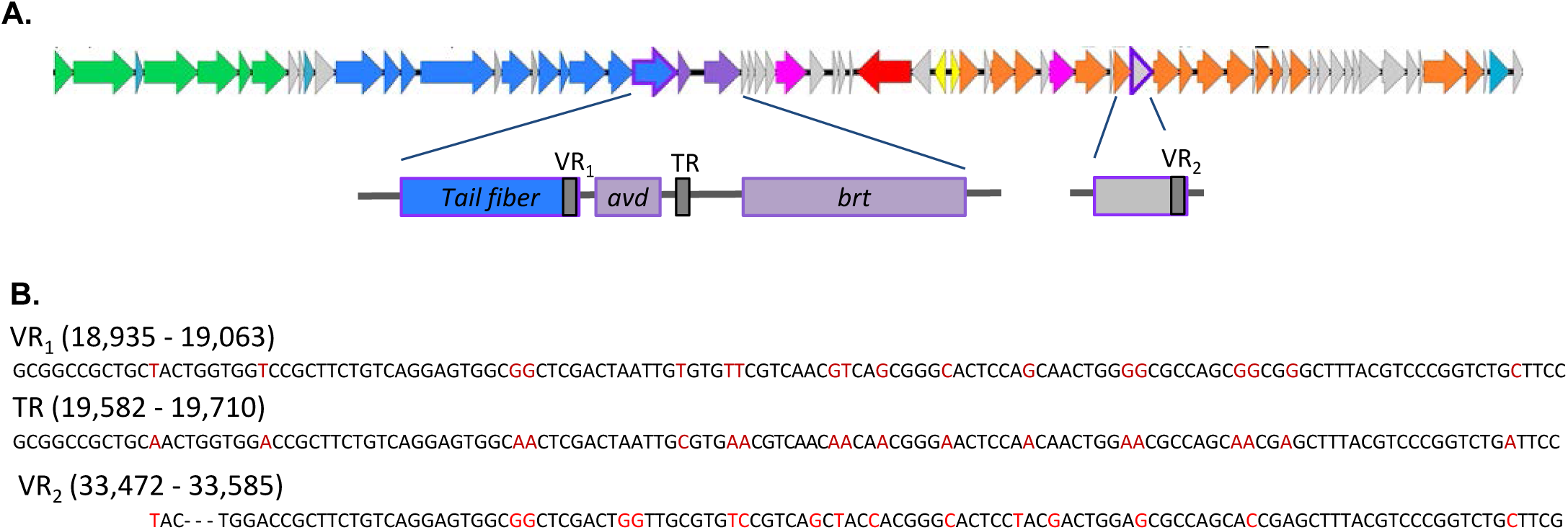
Diversity Generating Retroelement of Shimadzu. A) general organisation of the DGR in the Shimadzu genome. Color legend is as in Fig. 1. B) Sequences of the template repeat (TR), of the variable repeat VR_1_ in a tail gene (VR1), and of the second variable repeat VR_2_ in *orf44*. As already described, mainly TR adenines are modified in both VR (in red). Their positions in the Shimadzu genome are indicated (free phage form of the genome).

**Figure S3:**
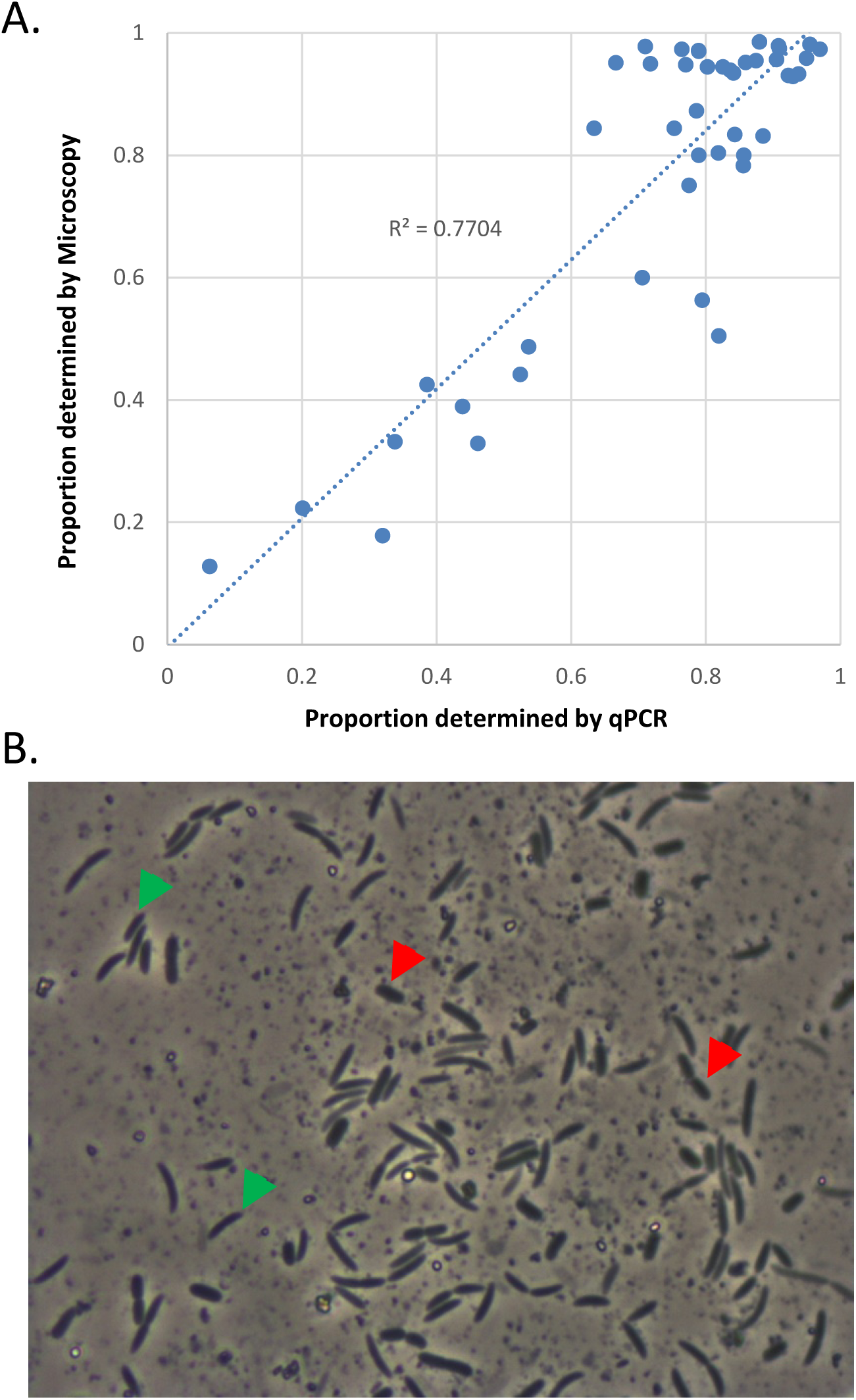
Comparison of quantification by qPCR and by microscopy. **A)** Proportion of *R. intestinalis* determined by quantitative PCR (qPCR) in function of the proportion determined by microscopy. An overall good correlation is observed. **B)** Observation of faeces by microscopy. *R. intestinalis* cells (green arrows) are clearly distinguishable from *E. coli* cells (red arrows).

**Figure S4:**
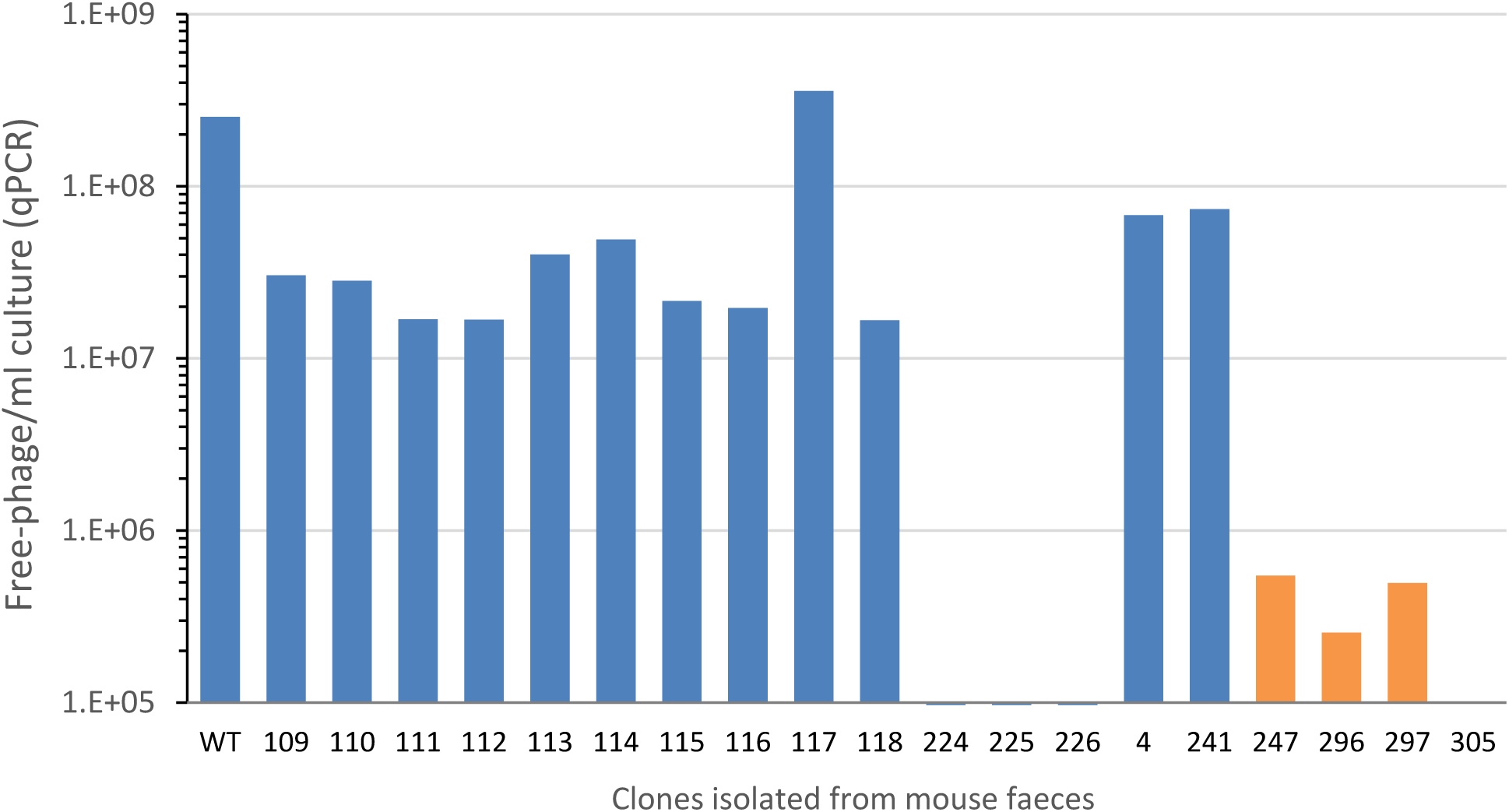
*In vitro* free phage production of clones isolated from mouse faeces. Shimadzu genome copy number in 16 hours saturated cultures. Clone numbers are those of Fig. 5. Orange bars correspond to clones in which a Shim prophage was detected. Clones 224, 225 226 and 305 are devoid of a Shimadzu prophage.

